# Age-varying DNA methylation patterns associated with blood pressure across the lifespan

**DOI:** 10.64898/2025.12.09.693332

**Authors:** Gang Xu, Xiaowei Zhuang, Amei Amei, Zuoheng Wang, Edwin C. Oh

## Abstract

**Background:** Epigenome-wide association studies (EWAS) have identified associations between DNA methylation and blood pressure, yet most rely on single-time-point data and cannot capture how methylation and blood pressure relationships change with age.

**Methods:** We conducted a longitudinal EWAS of 1,945 blood samples from 976 participants in the Multi-Ethnic Study of Atherosclerosis using a spline-based varying-coefficient model to detect age-dependent associations between DNA methylation in blood and blood pressure traits. Findings were evaluated for replication in 1,187 samples from the Framingham Heart Study. Models were adjusted for sex, ancestry, and leukocyte composition to account for cellular heterogeneity.

**Results:** Six CpG sites showed significant age-dependent associations with systolic or pulse pressure after correction for multiple testing. These included loci within *STIP1*, *CSRP1*, and *KDM6A* that replicated in the Framingham cohort. Several CpG sites demonstrated a reversal of effect direction with advancing age, where higher methylation was associated with higher systolic pressure in younger adults but lower pressure later in life. Pathway enrichment analyses identified focal adhesion, actin cytoskeleton remodeling, and Wnt/β-catenin signaling, which are processes relevant to vascular aging. Drug target mapping identified 23 FDA-approved agents interacting with genes at these loci.

**Conclusions:** Blood-derived DNA methylation shows dynamic age-related associations with blood pressure that likely reflect systemic or vascular aging processes rather than direct cellular mediation. Longitudinal analytical frameworks can reveal temporal patterns in epigenetic variation that are not detectable in single time point studies and may inform the discovery of biomarkers for age related cardiovascular risk.

## INTRODUCTION

Hypertension is a chronic condition characterized by persistently high blood pressure and serves as one of the primary preventable risk factors for cardiovascular disease and dementia^1–3^. Although the genetic and environmental factors that contribute to hypertension have been extensively studied, growing evidence indicates that DNA methylation, an epigenetic process that modifies gene expression without altering DNA sequence, may reflect how genetic susceptibility and environmental exposures jointly influence vascular health ^4–6^. Current insights largely stem from cross-sectional genetic and epigenetic studies, revealing significant methylation changes associated with vascular aging and cardiovascular risk^5,6^. However, these studies only capture static snapshots of methylation at single time points, potentially missing key temporal dynamics. Understanding how DNA methylation directly influences the progression of hypertension over the lifespan could therefore significantly enhance our mechanistic understanding, uncovering novel biomarkers and therapeutic targets that evolve with age.

Conventional epigenome-wide association studies (EWAS) typically employ analytical approaches, such as generalized estimating equations (GEE) and generalized linear mixed models (GLMM), that inherently assume methylation effects remain fixed throughout life. This assumption may overlook biologically critical temporal variations and dynamic gene-environment interactions, limiting the detection of age-specific epigenetic modifications crucial for understanding hypertension progression^7,8^. Recent advances in statistical modeling allow the evaluation of dynamic genomic and epigenetic effects over time. One such method, the Varying coefficient Mixed Model Association Test (VMMAT), employs spline-based modeling to effectively capture smooth temporal variations in dynamic genomic effects across age intervals^9–11^. Other longitudinal methods, including functional principal component analysis (FPCA), Bayesian hierarchical modeling, and penalized spline regression, have also shown promise for identifying complex age dependent molecular patterns. ^12–14^. Incorporating such dynamic longitudinal techniques into EWAS analyses could substantially improve our understanding of epigenetic regulation over the lifespan, identifying critical windows of susceptibility and guiding targeted therapeutic strategies for hypertension.

DNA methylation influences hypertension progression by modulating both established physiological mechanisms and emerging epigenetic pathways. Hypertension involves well-characterized biological pathways, such as the renin-angiotensin-aldosterone system (RAAS), renal sodium handling, and vascular endothelial function, all crucial in regulating blood pressure homeostasis^15–17^. Consistent methylation changes have been identified in key genes within these canonical pathways, including angiotensin-converting enzyme (ACE), aldosterone synthase (CYP11B2), and endothelial nitric oxide synthase (eNOS), further implicating epigenetic mechanisms in hypertension pathogenesis^18–20^. Additionally, recent data have highlighted novel methylation-driven regulatory processes influencing blood pressure through pathways previously underappreciated, such as genes involved in cytoskeletal remodeling, stress response signaling, inflammatory regulation, and epigenetic modulators themselves^6,21,22^. These emerging findings demonstrate how environmental exposures—including diet, psychosocial stress, physical activity, and lifestyle behaviors—can trigger methylation changes across both classical and newly identified pathways, thereby shaping individual susceptibility to hypertension throughout the life course. Integrating these diverse physiological and epigenetic perspectives could reveal critical mechanisms and potential intervention points, improving clinical strategies for hypertension prevention and management.

In this study, we applied the VMMAT approach to identify age-dependent associations between DNA methylation and blood pressure, addressing limitations of traditional cross-sectional EWAS methods. To our knowledge, this represents the first longitudinal EWAS investigating DNA methylation dynamics in hypertension. Using longitudinal data from the Multi-Ethnic Study of Atherosclerosis (MESA)^23^—a diverse cohort with comprehensive repeated measurements of blood pressure and DNA methylation—we identified significant age-dependent epigenetic associations with systolic blood pressure (SBP) and pulse pressure (PP). These associations were subsequently validated in an independent cohort, the Framingham Heart Study (FHS)^24,25^. Pathway analyses contextualized the identified methylation sites within biological networks, highlighting key cardiovascular-related pathways, such as focal adhesion, actin cytoskeletal remodeling, and catenin signaling. Integration with publicly available drug and gene databases further highlighted the potential therapeutic relevance of several identified loci. Together, these results demonstrate that longitudinal modeling can uncover dynamic age related associations between DNA methylation and blood pressure that are not captured by cross-sectional approaches and may reflect molecular features of vascular aging.

## RESULTS

### Study datasets

Our longitudinal EWAS utilized genome-wide DNA methylation data from two independent cohorts. The discovery dataset comprised 1,945 blood samples collected longitudinally from 976 participants across two examination periods within the Multi-Ethnic Study of Atherosclerosis (MESA). The replication dataset consisted of 1,187 blood samples from 1,187 independent participants in the Framingham Heart Study (FHS). Detailed demographic and clinical characteristics comparing participants from the MESA and FHS cohorts are summarized in **Table 1**.

**Table 1.** Demographic and clinical characteristics of study participants from the Multi-Ethnic Study of Atherosclerosis (MESA) and Framingham Heart Study (FHS).

| Characteristic | MESA | FHS |
| --- | --- | --- |
|  | *n* = 975 | *n* = 1,187 |
| Age at baseline (years), mean (SD) | 60.34 (9.66) | 72.11 (8.08) |
| Male, *n* (%) | 461 (47.28) | 562 (45.66) |
| SBP at baseline (years), mean (SD) | 122.89 (20.41) | 126.78 (16.55) |
| DBP at baseline (years), mean (SD) | 71.52 (10.16) | 70.68 (9.58) |
| PP at baseline (years), mean (SD) | 51.36 (15.95) | 56.10 (15.40) |

MESA: Multi-Ethnic Study of Atherosclerosis; FHS: Framingham Heart Study; SD: standard deviation; SBP: systolic blood pressure; DBP: diastolic blood pressure; PP: pulse pressure.

### Detection of age-varying methylation effects on blood pressure

Dynamic DNA methylation effects on blood pressure traits were assessed using the varying coefficient mixed model association test (VMMAT)^11^. Specifically, we evaluated associations with systolic blood pressure (SBP), diastolic blood pressure (DBP), and pulse pressure (PP), employing the joint Cauchy combination test^26^ implemented in VMMAT (see details in Methods). As a benchmark comparison, we also implemented a generalized linear mixed model (GLMM) assuming a static (age-invariant) methylation effect. Both models incorporated age (VMMAT use cubic splines to model age-varying intercept and methylation effect), sex, the first four genomic principal components (PCs) of ancestry^27^, and estimated cellular proportions (CD4T, NK, B cell, Mono, and Gran, excluding CD8T to mitigate multicollinearity^28^). Subject-specific random intercepts and slopes were included to capture within-individual correlations across repeated measures. The overall analytical workflow is depicted in **Figure 1**.

**Figure 1.**
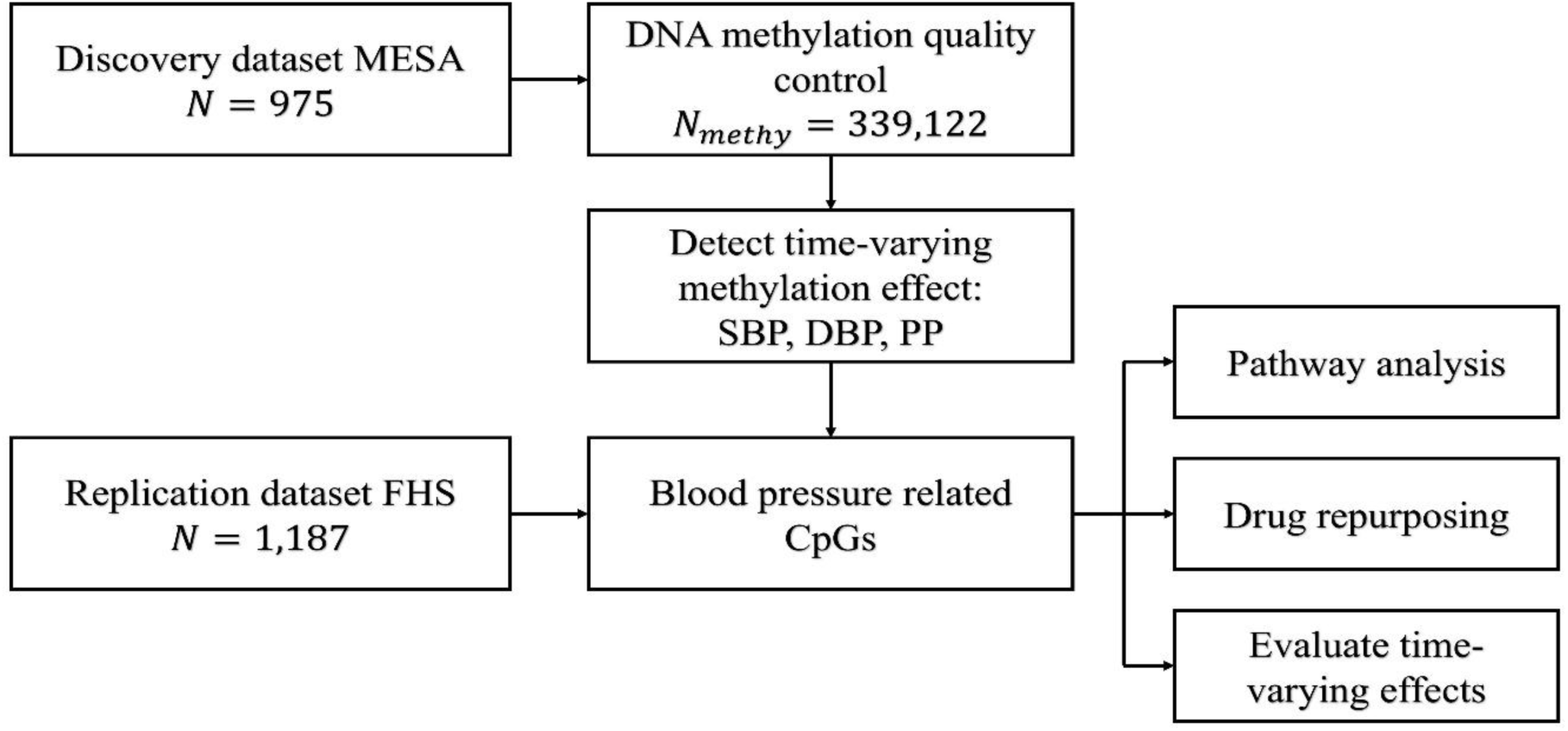
Analytical pipeline of the longitudinal methylation study. DNA methylation sites associated with blood pressure were identified using the varying coefficient mixed model association test (VMMAT) in the Multi-Ethnic Study of Atherosclerosis (MESA). Significant CpG sites underwent subsequent analyses, including: (1) functional enrichment analyses to identify relevant biological pathways; (2) drug repurposing analyses for potential therapeutic relevance; and (3) evaluation of age-dependent methylation effects on systolic and pulse pressures. Replication analyses were performed using the independent Framingham Heart Study (FHS) dataset.

Genome control inflation factors^29^ (λ) were computed to assess potential systemic bias in our association test statistics. Ideally, λ should equal 1; values below 1.1 are typically considered acceptable in genome-wide analyses^30^, with modest inflation reflecting polygenicity rather than confounding. In our study, SBP analyses yielded λ = 1.10 (VMMAT) and 1.00 (GLMM); PP analyses had λ = 1.13 (VMMAT) and 1.06 (GLMM); and DBP analyses showed λ = 0.99 (VMMAT) and 0.85 (GLMM). Although the VMMAT λ for PP slightly exceeded the conventional threshold, it is still within a range unlikely to compromise result validity, as thresholds up to ∼1.2 are often tolerated in polygenic traits^31^.

Applying a stringent Bonferroni-corrected genome-wide significance threshold of 1.47×10⁻⁷ (0.05/339,122 CpGs), we identified two significant CpG sites with age-varying effects on blood pressure: cg08023851 within the *STIP1* gene (p=1.37×10⁻⁷) significantly associated with SBP, and cg02165526 within the *CSRP1* gene (p=1.39×10⁻⁷) significantly associated with PP. To explore additional methylation signals at a less conservative threshold, we applied the Benjamini–Hochberg false discovery rate (FDR) correction^32^, identifying four additional CpG sites significantly associated with SBP (**Table 2** and **Figure 2A**). Despite evaluating associations with DBP, no CpGs reached genome-wide significance. Consequently, further analyses focused solely on SBP and PP, for which significant age-varying effects were evident.

**Figure 2.**
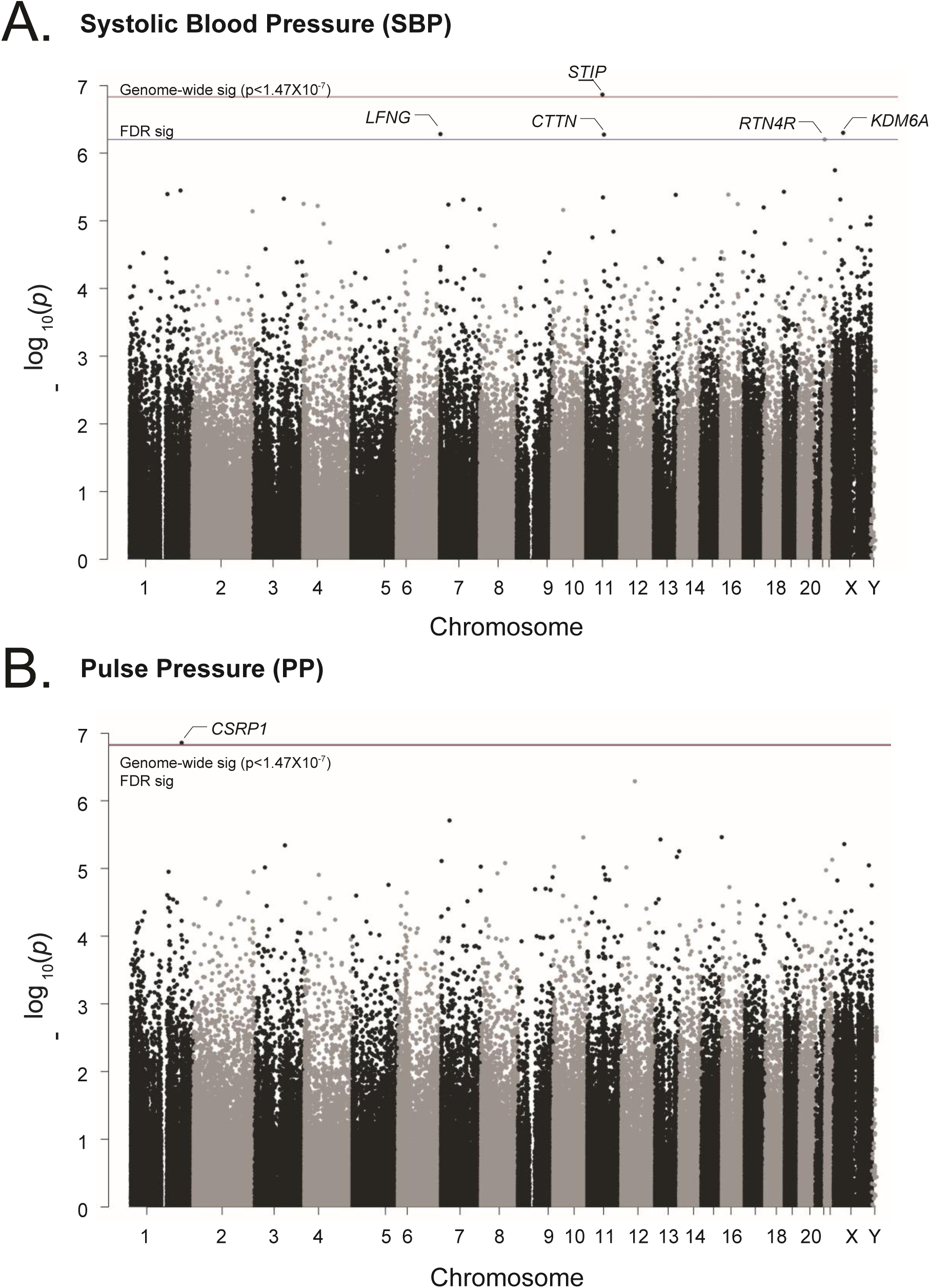
Genome-wide association analysis of methylation effects on blood pressure. Manhattan plots illustrate significant CpG sites associated with (A) systolic blood pressure (SBP) and (B) pulse pressure (PP). Genome-wide significant thresholds (Bonferroni-corrected, p<1.47×10⁻⁷) and False Discovery Rate (FDR)-corrected significant sites are indicated.

**Table 2.** Genome-wide significant CpG sites associated with blood pressure traits identified in the Multi-Ethnic Study of Atherosclerosis (MESA) and replicated in the Framingham Heart Study (FHS).

| Phenotype | Chr | Gene Region | CpG | Position | Discovery VMMAT | Discovery VMMAT_FDR | Discovery GLMM | Replication VMMAT |
| --- | --- | --- | --- | --- | --- | --- | --- | --- |
| SBP | 7 | LFNG | cg08196267 | 2,559,471 | 5.24E-07 | 0.04 | 1.50E-03 | 1.59E-01 |
|  | 11 | STIP1 | cg08023851 | 63,953,193 | **1.37E-07** | 0.04 | 3.50E-06 | **2.17E-06** |
|  | 11 | CTTN | cg27243121 | 70,281,584 | 5.31E-07 | 0.04 | 3.83E-04 | 2.15E-02 |
|  | 22 | RTN4R | cg03475420 | 20,255,667 | 6.28E-07 | 0.04 | 2.41E-04 | 1.54E-01 |
|  | X | KDM6A | cg05550588 | 44,732,733 | 5.00E-07 | 0.04 | 2.12E-01 | 6.76E-01 |
| PP | 1 | CSRP1 | cg02165526 | 201,478,358 | **1.39E-07** | 0.05 | 7.46E-06 | **1.41E-02** |

Discovery p-values are bold if they are smaller than 0.05/339,122 CpGs. Replication p-values are bold if they are smaller than 0.05/5 discovery CpGs for SBP and 0.05/1 discovery CpGs for PP.

Replication analyses in the independent FHS cohort validated our findings. Specifically, CpG cg08023851 (STIP1) demonstrated significant replication for SBP (p=2.17×10⁻⁶), surpassing the Bonferroni-corrected threshold (0.05/5 CpGs tested). Similarly, CpG cg02165526 (CSRP1) significantly replicated for PP (p=1.42×10⁻²). These robust replications underscore the reliability and generalizability of our primary findings.

Several significant CpGs mapped to genes previously implicated in blood pressure regulation. Specifically, *KDM6A* was associated with elevated blood pressure in mouse models^33^, *STIP1* variants were recently linked to DBP in a large-scale genome-wide association study (GWAS)^34^, *RTN4R* polymorphisms correlated with elevated systolic and diastolic pressures in the DASH trial^35^, and *LFNG* was identified as a locus for SBP in recent GWAS studies (**Table 2**)^36^.

### Age-varying methylation effects

We next evaluated the dynamic changes in methylation effects across different age groups for six significant CpGs identified in **Table 2**. The five CpGs associated with SBP demonstrated a progressive decrease in effect size and further changed in direction with advancing age, indicating that higher methylation at these loci correlated positively with systolic pressure in younger adults, transitioning to a negative correlation in older age groups (**Figure 3A**). The CpG associated with PP, cg02165526, on the other hand, exhibited a progressively stronger positive association between methylation and increased pulse pressure across age strata (**Figure 3B**). Collectively, these findings underscore the complex and dynamic age-dependent nature of DNA methylation’s role in blood-pressure regulation.

**Figure 3.**
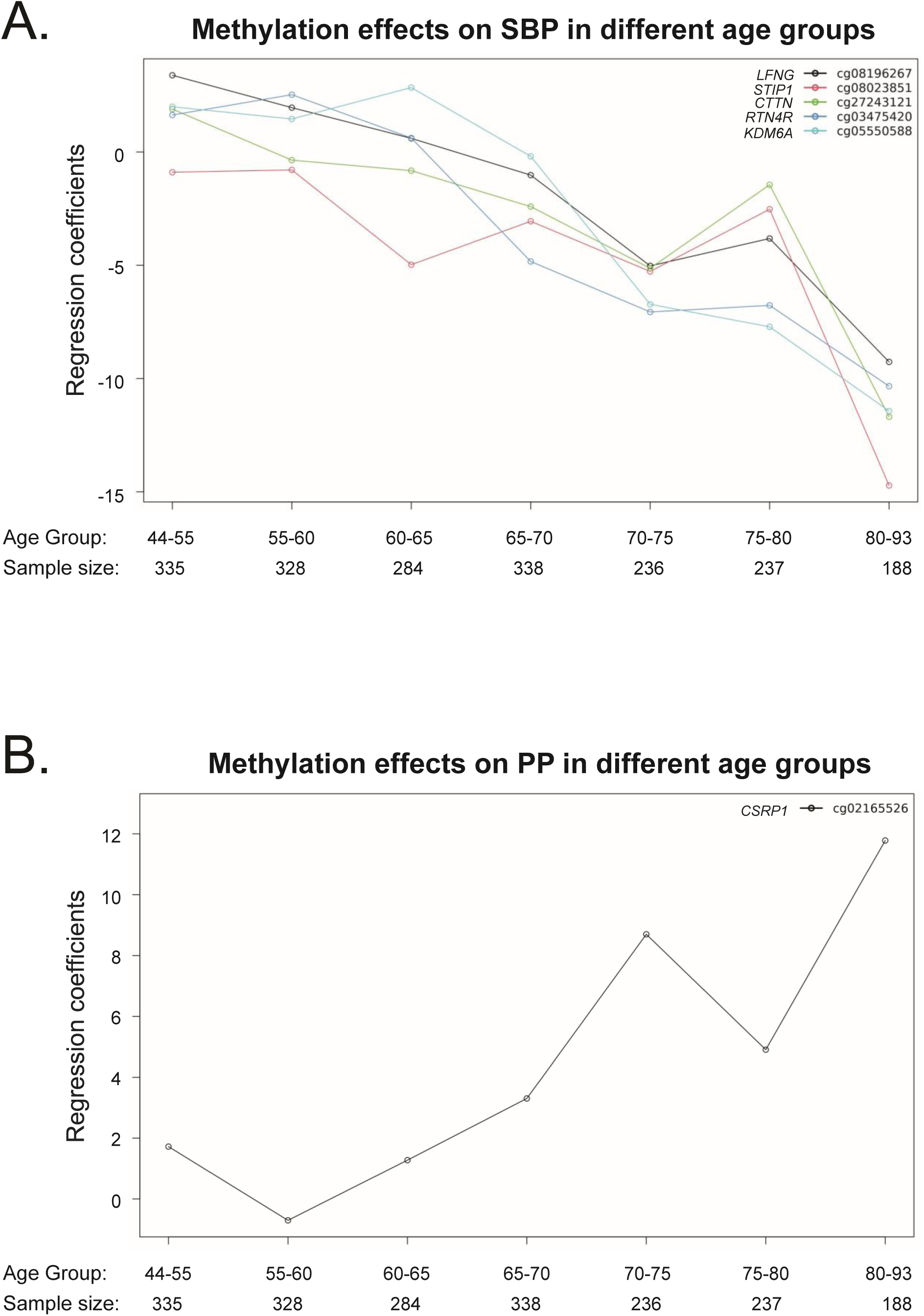
Age-dependent methylation effects on blood pressure. Linear regression coefficients illustrating the strength and direction of methylation effects on (A) systolic blood pressure (SBP) and (B) pulse pressure (PP) across different age groups. Each CpG is represented separately to highlight dynamic shifts in methylation impact across the lifespan.

### Pathway analysis

To contextualize the identified CpG associations biologically, we selected CpGs with p-values less than 5×10⁻⁴ from either VMMAT or GLMM analyses and mapped them to genes when CpGs were located within annotated genes. This selection yielded 521 genes for SBP and 586 genes for PP (**Supplementary Table 1 and 2)**, which were subsequently analyzed using PANTHER^37^ for functional pathway enrichment. Utilizing Gene Ontology (GO) terms—including biological processes, molecular functions, and cellular components—we identified three significantly enriched pathways related to blood pressure (FDR-p<0.05 in Fisher’s z test)^38^ (**Table 3, Supplementary Table 4 and 5**). The first significant pathway was focal adhesion (SBP: p=2.59×10⁻⁵, FDR-p=3.04×10⁻³; PP: p=2.95×10⁻⁶, FDR-p=7.37×10⁻⁴). Focal adhesion kinase (FAK) activation is known to influence vascular smooth muscle contraction and arterial stiffness, essential contributors to hypertension pathogenesis^39–41^. The second enriched pathway was actin cytoskeleton remodeling (PP: p=7.59×10⁻⁶, FDR-p=1.38×10⁻³), a pathway repeatedly implicated in both pulmonary and systemic hypertension due to its critical role in vascular tone regulation^42,43^. Lastly, we observed enrichment of the catenin complex pathway (SBP: p=7.29×10⁻⁴, FDR-p=4.28×10⁻²), specifically involving the Wnt/β-catenin signaling cascade, which modulates blood pressure in animal models and is actively investigated for its therapeutic potential in hypertension management^44,45^.

**Table 3.** Blood pressure related pathways based on the two gene lists linked to CpG sites associated with systolic blood pressure (SBP) or pulse pressure (PP).

| Phenotype | Pathways | P-values | FDR-p |
| --- | --- | --- | --- |
| SBP | focal adhesion | 2.59E-05 | 3.04E-03 |
|  | endosome | 2.87E-05 | 3.18E-03 |
|  | glutamatergic synapse | 1.48E-04 | 1.41E-02 |
|  | catenin complex | 7.29E-04 | 4.28E-02 |
| PP | focal adhesion | 2.95E-06 | 7.37E-04 |
|  | actin binding | 3.21E-06 | 3.28E-03 |
|  | myofibril | 7.21E-06 | 1.44E-03 |
|  | actin cytoskeleton | 7.59E-06 | 1.38E-03 |
|  | contractile muscle fiber | 1.31E-05 | 1.87E-03 |
|  | sarcomere | 8.49E-05 | 6.78E-03 |
|  | cation channel complex | 4.45E-04 | 2.61E-02 |
|  | calcium channel complex | 6.06E-04 | 3.18E-02 |

### Drug repurposing analysis

To identify therapeutic opportunities, we performed drug repurposing analyses by intersecting the 521 genes associated with SBP and the 586 genes associated with PP (CpG p-values < 5×10^−4^, same as above pathway analysis), resulting in 273 shared genes (**Supplementary Table 3**). Querying the DrugBank database (v5.1.12)^46^ for these overlapping genes yielded 23 FDA-approved drugs targeting at least one gene (**Table 4**). Among these, salmon calcitonin, though lacking clinical antihypertensive applications, increases blood pressure via the renin-angiotensin system in animal models^47^. Additionally, protamine sulfate, clinically used to counteract heparin anticoagulation, has demonstrated rapid blood pressure lowering through nitric oxide-mediated sympathetic inhibition and vasodilation^48,49^. Furthermore, antithrombin alfa, used clinically in thrombotic conditions, can prevent endotoxin-induced hypotension in animal models through suppression of inducible nitric oxide synthase^50^, highlighting potential relevance for clinical blood-pressure modulation.

**Table 4.**
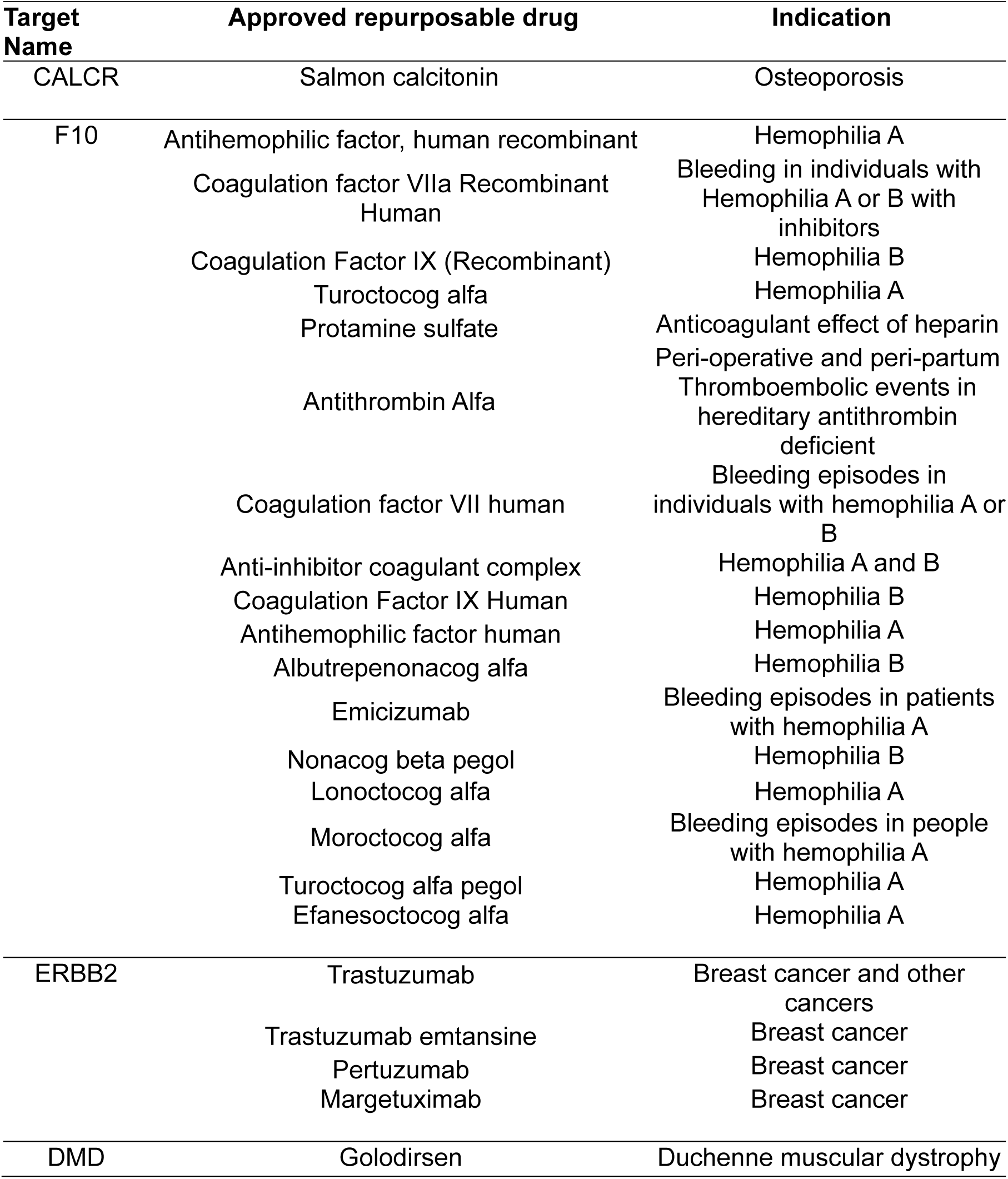
Drug repurposing analysis identifying FDA-approved drugs targeting genes linked to CpG sites associated with both systolic blood pressure (SBP) and pulse pressure (PP).

## DISCUSSION

Our longitudinal epigenome-wide association study (EWAS) identified several novel DNA methylation sites associated with blood pressure (BP), specifically highlighting significant CpG loci near the *STIP1*, *CSRP1*, *LFNG*, *KDM6A*, *RTN4R*, and *CTTN* genes. These loci exhibited genome-wide significant dynamic, age-dependent associations, where higher methylation was correlated with lower systolic blood pressure (SBP) in younger adults, but this relationship reversed in older individuals. Additionally, we observed a progressively stronger positive association between methylation and pulse pressure (PP) with advancing age, indicative of increased arterial stiffness. Pathway analyses further implicated focal adhesion, actin cytoskeletal remodeling, and Wnt/β-catenin signaling pathways, reinforcing their central roles in vascular aging and hypertension pathogenesis. Importantly, these associations were robustly replicated in an independent cohort from the Framingham Heart Study (FHS), underscoring their reliability and broader relevance.

A key innovation of our study was the development and application of a longitudinal design coupled with the Varying coefficient Mixed Model Association Test (VMMAT), providing insights inaccessible through conventional cross-sectional approaches. Previous large-scale meta-EWAS, including the CHARGE consortium^6^ and a subsequent EWAS involving ∼4,820 individuals of European and African ancestries^51^, identified several CpGs associated with blood pressure, predominantly within metabolic and inflammatory pathways. These studies demonstrated substantial overlap in CpG findings across populations, and validation in twin cohorts further reinforced shared genetic-epigenetic associations^51–53^. While our primary aim was to uncover dynamic, age-dependent methylation patterns, we noted partial overlap with previously reported loci, such as *PTPRN2*, *NR3C1*, *GABBR1*, and *FURIN*; however, these loci emerged as less significant findings in our analyses. For example, *PTPRN2* was identified in trans-ancestry DMR (Differentially Methylated Region) analyses of SBP^5^, and *NR3C1* methylation has been linked to blood pressure regulation in human cohort studies^54^. These earlier studies assumed constant methylation–phenotype relationships and did not account for temporal dynamics. In contrast, our VMMAT approach revealed CpG loci whose associations with blood pressure shift across the lifespan—suggesting protective effects early in life may become harmful later. This introduces a new dimension in epigenetic epidemiology, emphasizing that risk markers can exert age-dependent beneficial or harmful effects^6^. While some overlap in genes reinforces the general relevance of epigenetic mechanisms, our most significant findings uniquely emphasize the importance of age-dependent changes, thereby filling an essential knowledge gap and demonstrating that traditional cross-sectional methods may miss critical epigenetic indicators associated with hypertension.

The mechanistic implications of these age-varying methylation–BP links point to pathways of vascular aging and remodeling. Genes annotated to our top CpGs converge in pathways of focal adhesion, actin cytoskeleton dynamics, and catenin signaling, all fundamental to arterial structure and function^39,55,56^. For example, *CTTN* encodes cortactin, an actin-binding protein that stabilizes endothelial adherens junctions by linking VE-cadherin to the actin cytoskeleton through α/β-catenins^57,58^. Hypermethylation in the *CTTN* locus (likely reducing its expression) in younger individuals might enhance junctional stability and lower BP, whereas in older adults the same methylation could reflect or induce impaired repair of junctions under stress, contributing to higher BP. Similarly, *CSRP1* encodes a LIM-domain protein (CRP1) expressed in vascular smooth muscle; while not essential for baseline development, CRP1 modulates the smooth muscle response to arterial injury and stress^59,60^. The enrichment of Wnt/β-catenin signaling in our pathway analysis further supports a role in vascular aging; Wnt–catenin is a key driver of cell proliferation and senescence in the vasculature, and its dysregulation is linked to arterial calcification and stiffness in aging arteries^61,62^. Importantly, *KDM6A*, a histone demethylase identified here, has recently been implicated in blood pressure regulation and vascular integrity: loss of *KDM6A* in mice leads to increased arterial collagen deposition and stiffness^6^, raising BP, and KDM6A appears essential for normal renal sodium handling and BP homeostasis^33^. Our finding that higher methylation (presumably lower expression) of *KDM6A* is associated with rising BP in older adults is consistent with this, hinting that age-related epigenetic silencing of an epigenomic *regulator* might exacerbate vascular aging. In aggregate, these mechanistic links suggest that the CpGs we identified are not merely statistical associations but map to biological processes – cytoskeletal remodeling, cell–matrix adhesion, developmental signaling – that plausibly connect epigenetic drift to vascular aging and hypertension.

Despite its strengths, our study has important limitations. First, DNA methylation was measured in peripheral blood, which may not directly represent methylation changes in vascular tissues (endothelium or smooth muscle) that more directly govern BP. While some methylation signals are shared between blood and vascular tissue, others are tissue-specific; thus, our findings could be potential markers of systemic factors or immune processes. Future studies should examine these CpGs in vascular tissue or cell models to confirm functional relevance. Second, although longitudinal, our statistical model still test the association between BP and methylation rather than casual interpretation. It remains possible that changes in BP lead to altered methylation (rather than vice versa), or that a third factor (e.g., medication, diet, inflammation) influences both. We attempted to mitigate confounding by adjusting for key covariates, but residual confounding and reverse causation are challenging to eliminate which require additional biological meaningful assumptions. Third, our sample size, while sizeable for a longitudinal EWAS (nearly 1,000 participants, ∼1,945 samples), is modest by genome-wide standards, and we may have missed smaller effects. Moreover, multiple testing in EWAS is stringent, possibly omitting biologically relevant associations below significance thresholds. Nonetheless, replication of major findings in an independent cohort and pathway-level validations substantially mitigate these concerns. Additionally, our finding of no significant associations with diastolic blood pressure (DBP) may reflect distinct underlying epigenetic mechanisms for DBP, limitations in statistical power, or age-specific sensitivities, which should be explored further.

Looking forward, our study suggests several promising research directions. Future EWAS should routinely integrate longitudinal modeling to uncover dynamic epigenetic effects in other age-related phenotypes, such as glycemic control or cognitive decline. In hypertension specifically, integrating methylation data with transcriptomic^63^ and proteomic^64^ profiling could determine whether identified methylation shifts translate into gene expression or protein-level changes within vascular tissues. Such multi-omics approaches would greatly strengthen interpretations. Moreover, more frequent longitudinal sampling from younger adulthood through advanced ages could precisely pinpoint critical windows during which methylation exerts maximal influence on BP trajectories. Additionally, exploring relationships between BP-related methylation dynamics and accelerated biological aging (e.g., epigenetic clocks) may provide insights into how epigenetic markers predict cardiovascular risk over the life course^65–67^. Ultimately, our findings introduce a perspective where age-dependent epigenetic regulation dynamically influences cardiovascular trajectories, opening new avenues for targeted preventive and therapeutic strategies for hypertension management across the lifespan.

## Methods

### Study population of discovery dataset

Data for the discovery study were obtained from the public available dataset MESA from dbGaP (dbGaP Study Accession: phs000209). The MESA cohort comprises 6,814 asymptomatic adults aged 45–84 years, of whom 38% are White, 28% African American, 22% Hispanic, and 12% Asian—predominantly of Chinese ancestry^23^. The first blood examination spanned July 2000 – July 2002 and was followed by four additional examination visits.

### Study population of replication dataset

Data for the replication study were obtained from the public available dataset FHS from dbGaP (dbGaP Study Accession: phs000007). The FHS began in 1948 with 5,209 Framingham, Massachusetts residents aged 30–62 and has examined them every two years ever since^68^. In 1971 the study added 5,124 adult children and spouses (the Offspring cohort)^69^, and between 2002 and 2005 it enrolled 4,095 grandchildren (the Third Generation cohort)^70^, each cohort undergoing repeat exams roughly every four years.

### Ethics approval and consent

The MESA and FHS datasets were approved by their respective institutional review boards (IRBs). All participants provided written informed consent for participation and use of their genetic and clinical data. Our use of these publicly available datasets was reviewed and approved under an exempt status by the IRB at our institution (University of Nevada Las Vegas).

### DNA Methylation profile and quality control

DNA methylation was profiled in whole blood with the Illumina’s Methylation EPIC BeadChip (850 K) for 976 MESA participants who provided samples at Exams 1 and 5 as part of the National Heart, Lung, and Blood Institute (NHLBI) Trans-Omics for Precision Medicine (TOPMed) MESA Multi-Omics project. We performed DNA methylation quality control with the R package minfi^71^. CpG probes with detection p-values > 0.01 were removed, yielding 339,122 high-quality sites. The derived beta (𝛽) values were logit transformed to M values for the downstream EWAS analysis.

For replication, whole-blood DNA methylation in the Framingham Heart Study (FHS) was assayed with the Illumina Infinium Methylation EPIC BeadChip (850 K sites) in 1,249 Offspring-cohort participants and 547 Generation 3 participants enrolled in the NHLBI TOPMed FHS Multi-Omics project. Downstream analyses were restricted to the 1,249 Offspring participants, for whom we extracted M-values from the processed methylation data.

Cell type compositions ( CD8T, CD4T, NK, Bcell, monocyte and granulocyte) were estimated in both MESA and FHS with the estimateCellCounts function in R package minfi^71^ at default settings. Cell type composition estimators for CD4T, NK, Bcell, monocyte and granulocyte were then included as covariates in all EWAS models.

### Statistical modeling for age-varying methylation effects

We modeled longitudinal outcomes using a generalized linear mixed model (GLMM) with varying coefficients allowing for methylation effects to vary smoothly over age. Specifically, let 𝑌_𝑖,𝑗_ denote the trait value measured for subject 𝑖 (𝑖 = 1, … , 𝑛) at age 𝑡_𝑖𝑗_ (𝑗 = 1, … , 𝑚_𝑖_). Suppose a trait is measured repeatedly on a sample of 𝑛 subjects. For each observation we have a 𝑝-dimensional covariate vector 𝑿_𝑖𝑗_ which can include both static variables such as sex or dynamic variables such as cell type compositions and a DNA methylation measure 𝑮_𝑖𝑗_ expressed on the M value scale (−∞ < 𝑮_𝑖𝑗_ < ∞).

The model is defined as:

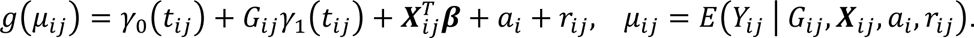

Here 𝑔(⋅) is the link function (logit for binary traits, identity for continuous traits). The functions 𝛾_0_(𝑡) and 𝛾_1_(𝑡) are smooth nonparametric functions of age 𝑡 representing a age-varying intercept and a age-varying methylation effect at the CpG under test, 𝜷 is the effects of the covariates. Correlation among repeat measurements is accommodated by two random terms: a subject-specific intercept 𝑎_𝑖_ ∼ 𝑁(0, 𝜎^2^) and a vector of time-dependent random effects 𝒓_𝑖_ = (𝑟_𝑖,1_, … , 𝑟_𝑖,𝑚_ ) ∼ 𝑀𝑉𝑁(𝟎, 𝜎^2^𝑹_𝑖_) with 𝑹_𝑖_ following an AR(1) structure parameterized by 𝜏 ^72,73^. Conditional on 𝑎_𝑖_ and 𝒓_𝑖_, the responses 𝑌_𝑖𝑗_ are assumed to be independent.

The primary hypothesis of interest is whether there is a age-varying methylation effect between the methylation site and the trait. We test

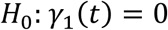

using the VMMAT ^11^, which combines the p-values of the spline basis coefficient and the variance component into a single Cauchy test statistic and provides a computationally efficient genome-wide screen for CpG sites with age-dependent associations.

### Pathway analysis

To investigate the biological functions of CpG sites associated with blood pressure, we conducted functional pathway enrichment using the PANTHER Classification System^37^. CpGs with p-values <5×10⁻⁴ from either VMMAT or GLMM analyses were mapped to genes located within annotated regions, resulting in 521 genes for SBP and 586 for PP (**Supplementary Tables 1–2**). Over-representation of Gene Ontology (GO) terms was evaluated using Fisher’s exact test, and statistical significance was controlled via Benjamini–Hochberg FDR (threshold: FDR-p<0.05).

### Drug repurposing analysis

To identify potential therapeutics, we focused on 273 genes associated with both SBP and PP (CpGs p < 5×10⁻⁴; **Supplementary Table 3**). These shared genes were cross-referenced with the DrugBank database (v5.1.12)^46^ to identify FDA-approved drugs targeting them. DrugBank provides comprehensive information on drug–target interactions, enabling systematic repurposing opportunities.

### Age-varying effects of significant CpG sites

To examine the temporal dynamics in methylation effects, we stratified the cohort into seven sequential age groups. Within each stratum, we refitted regression models to estimate age-specific methylation coefficients for each significant CpG site (as listed in **Table 2**). Coefficients were then plotted across age strata with lines connecting adjacent age groups to visualize trajectories of methylation effects across the lifespan.

## Declarations Ethical approval

This study involves secondary analyses of existing, de-identified data from the Multi-Ethnic Study of Atherosclerosis (MESA) and the Framingham Heart Study (FHS). All original cohort protocols were approved by the local Institutional Review Boards or Ethics Committees at each participating institution, and all participants provided written informed consent at enrollment. The present secondary analysis was reviewed and approved by the Human Research Protection Program Institutional Review Board at Yale University (IRB protocol 2000036485; expedited review; approval date November 21, 2023) and by the Biomedical Institutional Review Board at the University of Nevada, Las Vegas (UNLV-2025-267, “Secondary multi-omics data analyses to understand genetic effect of Hypertension”; expedited review; approval date July 24, 2025). All procedures were conducted in accordance with the Declaration of Helsinki, the U.S. Federal Policy for the Protection of Human Subjects (45 CFR 46), and relevant institutional guidelines and regulations.

## Availability of data and materials

The DNA-methylation and phenotypic data analyzed in this study were obtained from dbGaP under the following study accessions: MESA cohort data, phs000209; MESA TOPMed DNA methylation, phs001416; FHS cohort data, phs000007; and FHS TOPMed DNA methylation, phs000974. Any request to access relevant data in this study should be sent to the corresponding authors.

## Acknowledgments

The Multi-Ethnic Study of Atherosclerosis is supported by contracts 75N92020D00001, HHSN268201500003I, N01-HC-95159, 75N92020D00005, N01-HC-95160, 75N92020D00002, N01-HC-95161, 75N92020D00003, N01-HC-95162, 75N92020D00006, N01-HC-95163, 75N92020D00004, N01-HC-95164, 75N92020D00007, N01-HC-95165, N01-HC-95166, N01-HC-95167, N01-HC-95168 and N01-HC-95169 from the National Heart, Lung, and Blood Institute, and by grants UL1-TR-000040, UL1-TR-001079, and UL1-TR-001420 from the National Center for Advancing Translational Sciences (NCATS). The authors thank the other investigators, the staff, and the participants of the MESA study for their valuable contributions. A full list of participating MESA investigators and institutions can be found at http://www.mesa-nhlbi.org. This manuscript was not prepared in collaboration with investigators of the MESA and does not necessarily reflect the opinions or conclusions of the MESA or the NHLBI.

The Framingham Heart Study is conducted and supported by the National Heart, Lung, and Blood Institute (NHLBI) in collaboration with Boston University (Contract No. N01-HC-25195, HHSN268201500001I and 75N92019D00031). This manuscript was not prepared in collaboration with investigators of the Framingham Heart Study and does not necessarily reflect the opinions or views of the Framingham Heart Study, Boston University, or NHLBI.

## Author contributions

GX, XZ, AA, and EO contributed to the study design and interpretation of the data; GX and XZ collected, curated, and visualized the data; GX, AA, and ZW designed the statistical analyses; GX, XZ, and EO analyzed the data; GX and EO drafted the manuscript. All authors contributed to the article and approved the submitted version.

## REFERENCES

1 Campbell, N. R. et al. High blood pressure 2016: Why prevention and control are urgent and important. The world hypertension league, international society of hypertension, world stroke organization, international diabetes foundation, international council of cardiovascular prevention and rehabilitation, international society of nephrology. The Journal of Clinical Hypertension 18, 714 (2016).

2 Fuchs, F. D. & Whelton, P. K. High Blood Pressure and Cardiovascular Disease. Hypertension 75, 285–292 (2020). 10.1161/HYPERTENSIONAHA.119.14240

3 Ou, Y. N. et al. Blood Pressure and Risks of Cognitive Impairment and Dementia: A Systematic Review and Meta-Analysis of 209 Prospective Studies. Hypertension 76, 217–225 (2020). 10.1161/HYPERTENSIONAHA.120.14993

4 Karabaeva, R. Z. et al. Epigenetics of hypertension as a risk factor for the development of coronary artery disease in type 2 diabetes mellitus. Frontiers in Endocrinology 15, 1365738 (2024).

5 Kazmi, N. et al. Associations between high blood pressure and DNA methylation. PloS one 15, e0227728 (2020).

6 Richard, M. A. et al. DNA Methylation Analysis Identifies Loci for Blood Pressure Regulation. Am J Hum Genet 101, 888–902 (2017). 10.1016/j.ajhg.2017.09.028

7 Sitlani, C. M. et al. Generalized estimating equations for genome-wide association studies using longitudinal phenotype data. Statistics in medicine 34, 118–130 (2015). 10.1002/sim.6323

8 Nustad, H. E. et al. Epigenetics, heritability and longitudinal analysis. BMC Genet 19, 77 (2018). 10.1186/s12863-018-0648-1

9 Gorczak, K., Burzykowski, T. & Claesen, J. A varying-coefficient model for the analysis of methylation sequencing data. Comput Biol Chem 111, 108094 (2024). 10.1016/j.compbiolchem.2024.108094

10 Chen, H. & Wang, Y. A penalized spline approach to functional mixed effects model analysis. Biometrics 67, 861–870 (2011). 10.1111/j.1541-0420.2010.01524.x

11 Xu, G. et al. Retrospective varying coefficient association analysis of longitudinal binary traits: application to the identification of genetic loci associated with hypertension. The Annals of Applied Statistics 18, 487–505 (2024).

12 Friedman, N. Inferring cellular networks using probabilistic graphical models. Science 303, 799–805 (2004). 10.1126/science.1094068

13 Crainiceanu, C. M., Ruppert, D. & Wand, M. P. Bayesian analysis for penalized spline regression using WinBUGS. Journal of Statistical Software 14 (2005).

14 Fong, S. et al. Principal component-based clinical aging clocks identify signatures of healthy aging and targets for clinical intervention. Nat Aging 4, 1137–1152 (2024). 10.1038/s43587-024-00646-8

15 Stadt, M. & Layton, A. T. Modulation of blood pressure by dietary potassium and sodium: sex differences and modeling analysis. Am J Physiol Renal Physiol 328, F406–F417 (2025). 10.1152/ajprenal.00222.2024

16 Strazzullo, P., Galletti, F. & Barba, G. Altered renal handling of sodium in human hypertension: short review of the evidence. Hypertension 41, 1000–1005 (2003). 10.1161/01.HYP.0000066844.63035.3A

17 Coffman, T. M. The inextricable role of the kidney in hypertension. J Clin Invest 124, 2341–2347 (2014). 10.1172/JCI72274

18 Takeda, Y., Demura, M., Yoneda, T. & Takeda, Y. Epigenetic Regulation of the Renin-Angiotensin-Aldosterone System in Hypertension. Int J Mol Sci 25 (2024). 10.3390/ijms25158099

19 Kawakami-Mori, F. et al. Aberrant DNA methylation of hypothalamic angiotensin receptor in prenatal programmed hypertension. JCI Insight 3 (2018). 10.1172/jci.insight.95625

20 Yoshii, Y. et al. Hypomethylation of CYP11B2 in Aldosterone-Producing Adenoma. Hypertension 68, 1432–1437 (2016). 10.1161/HYPERTENSIONAHA.116.08313

21 Mangum, K. D. et al. Epigenetic alteration of smooth muscle cells regulates endothelin-dependent blood pressure and hypertensive arterial remodeling. J Clin Invest 135 (2025). 10.1172/JCI186146

22 Chaudhary, M. Novel methylation mark and essential hypertension. J Genet Eng Biotechnol 20, 11 (2022). 10.1186/s43141-022-00301-y

23 Bild, D. E. et al. Multi-ethnic study of atherosclerosis: objectives and design. American journal of epidemiology 156, 871–881 (2002). 10.1093/aje/kwf113

24 Mahmood, S. S., Levy, D., Vasan, R. S. & Wang, T. J. The Framingham Heart Study and the epidemiology of cardiovascular disease: a historical perspective. Lancet 383, 999–1008 (2014). 10.1016/S0140-6736(13)61752-3

25 Tsao, C. W. & Vasan, R. S. Cohort Profile: The Framingham Heart Study (FHS): overview of milestones in cardiovascular epidemiology. Int J Epidemiol 44, 1800–1813 (2015). 10.1093/ije/dyv337

26 Liu, Y. & Xie, J. Cauchy combination test: a powerful test with analytic p-value calculation under arbitrary dependency structures. Journal of the American Statistical Association 115, 393–402 (2020).

27 Hu, X. et al. Multi-ancestry epigenome-wide analyses identify methylated sites associated with aortic augmentation index in TOPMed MESA. Scientific reports 13, 17680 (2023).

28 Vasanthakumar, A. et al. Harnessing peripheral DNA methylation differences in the Alzheimer’s Disease Neuroimaging Initiative (ADNI) to reveal novel biomarkers of disease. Clinical epigenetics 12, 1–11 (2020).

29 Yang, J. et al. Genomic inflation factors under polygenic inheritance. Eur J Hum Genet 19, 807–812 (2011). 10.1038/ejhg.2011.39

30 Williams, C. J. et al. Genome wide association study of response to interval and continuous exercise training: the Predict-HIIT study. Journal of biomedical science 28, 37 (2021).

31 Nabais, M. F. et al. Meta-analysis of genome-wide DNA methylation identifies shared associations across neurodegenerative disorders. Genome biology 22, 1–30 (2021).

32 Benjamini, Y. & Hochberg, Y. Controlling the false discovery rate: a practical and powerful approach to multiple testing. Journal of the Royal statistical society: series B (Methodological) 57, 289–300 (1995).

33 Han, X., Akinseye, L. & Sun, Z. KDM6A demethylase regulates renal sodium excretion and blood pressure. Hypertension 81, 541–551 (2024).

34 Shi, S. et al. A Genomics England haplotype reference panel and imputation of UK Biobank. Nature Genetics 56, 1800–1803 (2024).

35 Kim, H., Lichtenstein, A. H., Coresh, J., Appel, L. J. & Rebholz, C. M. Serum protein responses to Dietary Approaches to Stop Hypertension (DASH) and DASH-Sodium trials and associations with blood pressure changes. Journal of Hypertension 42, 1823–1830 (2024).

36 Koskeridis, F. et al. Multi-trait association analysis reveals shared genetic loci between Alzheimer’s disease and cardiovascular traits. Nature Communications 15, 9827 (2024).

37 Mi, H. et al. PANTHER version 16: a revised family classification, tree-based classification tool, enhancer regions and extensive API. Nucleic acids research 49, D394–D403 (2021).

38 Ashburner, M. et al. Gene ontology: tool for the unification of biology. Nature genetics 25, 25–29 (2000).

39 Ribeiro-Silva, J., Miyakawa, A. & Krieger, J. E. Focal adhesion signaling: vascular smooth muscle cell contractility beyond calcium mechanisms. Clinical Science 135, 1189–1207 (2021).

40 Sugimura, K. et al. Hypertension promotes phosphorylation of focal adhesion kinase and proline-rich tyrosine kinase 2 in rats: implication for the pathogenesis of hypertensive vascular disease. The Tohoku Journal of Experimental Medicine 222, 201–210 (2010).

41 O’Brien, H. E. et al. Expression quantitative trait loci in the developing human brain and their enrichment in neuropsychiatric disorders. Genome biology 19, 1–13 (2018).

42 Luo, J. et al. Integrative analyses of gene expression profile reveal potential crucial roles of mitotic cell cycle and microtubule cytoskeleton in pulmonary artery hypertension. BMC medical genomics 13, 1–14 (2020).

43 Paul, A. S. et al. FXR1 regulates vascular smooth muscle cell cytoskeleton, VSMC contractility, and blood pressure by multiple mechanisms. Cell reports 42 (2023).

44 Xiao, L. et al. Wnt/β-catenin regulates blood pressure and kidney injury in rats. Biochimica Et Biophysica Acta (BBA)-Molecular Basis of Disease 1865, 1313–1322 (2019).

45 Zhao, Y. et al. An essential role for Wnt/β-catenin signaling in mediating hypertensive heart disease. Scientific reports 8, 8996 (2018).

46 Wishart, D. S. et al. DrugBank: a knowledgebase for drugs, drug actions and drug targets. Nucleic acids research 36, D901–D906 (2008).

47 Clementi, G. et al. Effects of salmon calcitonin on plasma renin activity and systolic blood pressure in the rat. Neuroscience letters 66, 351–355 (1986).

48 Oe, M., Asou, T., Morita, S., Yasui, H. & Tokunaga, K. Protamine-induced hypotension in heart operations: application of the concept of ventricular-arterial coupling. The Journal of Thoracic and Cardiovascular Surgery 112, 462–471 (1996).

49 Hamada, Y., Kameyama, Y., Narita, H., Benson, K. T. & Goto, H. Protamine after heparin produces hypotension resulting from decreased sympathetic outflow secondary to increased nitric oxide in the central nervous system. Anesthesia & Analgesia 100, 33–37 (2005).

50 Isobe, H., Okajima, K., Uchiba, M., Harada, N. & Okabe, H. Antithrombin prevents endotoxin-induced hypotension by inhibiting the induction of nitric oxide synthase in rats. *Blood*, The Journal of the American Society of Hematology 99, 1638–1645 (2002).

51 Huang, Y. et al. Identification, Heritability, and Relation With Gene Expression of Novel DNA Methylation Loci for Blood Pressure. Hypertension 76, 195–205 (2020). 10.1161/HYPERTENSIONAHA.120.14973

52 Jhun, M. A. et al. A multi-ethnic epigenome-wide association study of leukocyte DNA methylation and blood lipids. Nat Commun 12, 3987 (2021). 10.1038/s41467-021-23899-y

53 Zhang, X. et al. The Interplay of Epigenetic, Genetic, and Traditional Risk Factors on Blood Pressure: Findings from the Health and Retirement Study. Genes (Basel) 13 (2022). 10.3390/genes13111959

54 Dwi Putra, S. E., et al. DNA methylation of the glucocorticoid receptor gene promoter in the placenta is associated with blood pressure regulation in human pregnancy. J Hypertens 35, 2276–2286 (2017). 10.1097/HJH.0000000000001450

55 Ziegler, N. et al. beta-Catenin Is Required for Endothelial Cyp1b1 Regulation Influencing Metabolic Barrier Function. J Neurosci 36, 8921–8935 (2016). 10.1523/JNEUROSCI.0148-16.2016

56 Ciobanasu, C., Faivre, B. & Le Clainche, C. Actin dynamics associated with focal adhesions. Int J Cell Biol 2012, 941292 (2012). 10.1155/2012/941292

57 Moztarzadeh, S. et al. Cortactin is in a complex with VE-cadherin and is required for endothelial adherens junction stability through Rap1/Rac1 activation. Sci Rep 14, 1218 (2024). 10.1038/s41598-024-51269-3

58 Abu Taha, A. & Schnittler, H. J. Dynamics between actin and the VE-cadherin/catenin complex: novel aspects of the ARP2/3 complex in regulation of endothelial junctions. Cell Adh Migr 8, 125–135 (2014). 10.4161/cam.28243

59 Lilly, B. et al. Loss of the serum response factor cofactor, cysteine-rich protein 1, attenuates neointima formation in the mouse. Arterioscler Thromb Vasc Biol 30, 694–701 (2010). 10.1161/ATVBAHA.109.200741

60 Chang, D. F. et al. Cysteine-rich LIM-only proteins CRP1 and CRP2 are potent smooth muscle differentiation cofactors. Dev Cell 4, 107–118 (2003). 10.1016/s1534-5807(02)00396-9

61 Bundy, K., Boone, J. & Simpson, C. L. Wnt Signaling in Vascular Calcification. Front Cardiovasc Med 8, 708470 (2021). 10.3389/fcvm.2021.708470

62 Vallee, A. Arterial Stiffness and the Canonical WNT/beta-catenin Pathway. Curr Hypertens Rep 24, 499–507 (2022). 10.1007/s11906-022-01211-7

63 Skovgaard, A. C., Nejad, A. M., Beck, H. C., Tan, Q. & Soerensen, M. Epigenomics and transcriptomics association study of blood pressure and incident diagnosis of hypertension in twins. Hypertens Res 48, 1599–1612 (2025). 10.1038/s41440-025-02164-5

64 Zaghlool, S. B. et al. Epigenetics meets proteomics in an epigenome-wide association study with circulating blood plasma protein traits. Nat Commun 11, 15 (2020). 10.1038/s41467-019-13831-w

65 Xiao, L. et al. Associations Between Blood Pressure and Accelerated DNA Methylation Aging. J Am Heart Assoc 11, e022257 (2022). 10.1161/JAHA.121.022257

66 Wang, X. M., Wang, H. P., Chen, H. Z. & Liu, D. P. Epigenetic Clock: Future of Hypertension Prediction? Hypertension 80, 1569–1571 (2023). 10.1161/HYPERTENSIONAHA.123.21197

67 Carbonneau, M. et al. Epigenetic Age Mediates the Association of Life’s Essential 8 With Cardiovascular Disease and Mortality. J Am Heart Assoc 13, e032743 (2024). 10.1161/JAHA.123.032743

68 Dawber, T. R., Meadors, G. F. & Moore Jr, F. E. Epidemiological approaches to heart disease: the Framingham Study. American Journal of Public Health and the Nations Health 41, 279–286 (1951).

69 Feinleib, M., Kannel, W. B., Garrison, R. J., McNamara, P. M. & Castelli, W. P. The Framingham offspring study. Design and preliminary data. Preventive medicine 4, 518–525 (1975).

70 Splansky, G. L. et al. The third generation cohort of the National Heart, Lung, and Blood Institute’s Framingham Heart Study: design, recruitment, and initial examination. American journal of epidemiology 165, 1328–1335 (2007).

71 Fortin, J.-P., Triche Jr, T. J. & Hansen, K. D. Preprocessing, normalization and integration of the Illumina HumanMethylationEPIC array with minfi. Bioinformatics 33, 558–560 (2017).

72 Wu, W. et al. Retrospective Association Analysis of Longitudinal Binary Traits Identifies Important Loci and Pathways in Cocaine Use. Genetics 213, 1225–1236 (2019). 10.1534/genetics.119.302598

73 Wang, Z., Xu, K., Zhang, X., Wu, X. & Wang, Z. Longitudinal SNP-set association analysis of quantitative phenotypes. Genetic epidemiology 41, 81–93 (2017).

